# Limitations of differential expression for cell-type marker discovery in single-cell RNA sequencing atlases

**DOI:** 10.64898/2026.09.25.754486

**Authors:** Kellen Doggett, Daniel Pintard, Richard H. Scheuermann, Yun Zhang

## Abstract

Cell-type marker genes are definitional characteristics of a cell type, and in single-cell RNA sequencing atlases they are commonly nominated through differential expression (DE) testing. The Wilcoxon rank-sum test is the default method for marker gene discovery in Seurat and a recommended method in Scanpy, two widely used computational software packages for scRNA-seq analysis. Structurally, Wilcoxon’s standardized statistic grows with sample size at a fixed underlying effect, meaning computed rankings across clusters of unequal size are incomparable. We quantified this effect across three atlas datasets - lung, kidney, and brain. Mean Wilcoxon score scaled with log cluster size at Pearson correlation *r* = 0.90-0.95 in each organ. In contrast, a classification-based comparator, NS-Forest, showed low correlation with cluster size (*r* = 0.10-0.28). This DE-cluster size relationship remained essentially unchanged after controlling for classification performance through partial correlation, decayed monotonically under cluster size downsampling, and was reproduced in simulation with marker quality fixed. The Wilcoxon rank-sum test can be transformed into a rank-based effect size by normalizing by the target-background pair count, removing the positive size dependence. However, this effect size nominates an identical set of candidate genes as the original statistic, indicating the cluster size dependence and poor specificity of DE nominated genes are separate problems. Genes nominated under default settings were substantially less specific to their target clusters, with median on-target fractions of 0.204 (lung), 0.166 (kidney), and 0.083 (brain), compared to NS-Forest’s marker genes (0.580, 0.687, 0.364), revealing the relatively poor specificity that highly scoring DE genes exhibit.

## 1 Introduction

Cell-type marker genes are definitional characteristics of a cell type (Bakken et al., 2017). Marker genes are the link between cell type populations identified computationally across datasets and their use in downstream experimental research. The quality of marker genes determines the accuracy with which a cell type can be identified outside of the original dataset in which it was first defined.

In modern single-cell RNA sequencing (scRNA-seq) atlases, markers are commonly identified through differential expression (DE) testing in a one-versus-rest format. The Wilcoxon rank-sum test is the default setting in Seurat’s *FindAllMarkers* and is a recommended method in Scanpy’s documentation (Hao et al., 2024; Wolf et al., 2018). Candidate genes are ranked by the returned statistic, and the top scoring genes are reported as the cluster’s markers.

There is a fundamental mismatch between what differential expression computes and what marker selection requires. The Wilcoxon rank-sum test asks whether two samples are drawn from the same distribution, whereas marker selection asks whether a gene can distinguish a cell type from all others. Wilcoxon is properly calibrated under unequal group sizes, but its standardized statistic is not an effect size, and its computed magnitude is not comparable across comparisons of different sizes. This is exacerbated by the heterogeneous population of cell types in scRNA-seq data. The test’s standardized statistic is structurally a power statistic; at a fixed underlying effect, its magnitude grows with the comparison group. The interaction between cluster size imbalance and marker ranking across atlases has yet to be quantified.

This report addresses three questions. First, is DE-based marker selection confounded by cluster size? Second, are the genes that DE nominates less specific to their target cluster than markers chosen by a structurally different method, one that maximizes classification power? Third, can either of these problems be controlled through a transformation of the Wilcoxon rank-sum formula?

As a comparator to DE at its default settings, we use NS-Forest v4.1 as described in Liu et al. (2024). NS-Forest optimizes its marker gene candidate set on classification F-beta score instead of a test statistic. Its selection criterion is computed through proportions and has no structural reason to scale with target cluster size. This makes NS-Forest an apt comparison against DE statistic’s behavior. An additional comparison group granting DE the opportunity to use an NS-Forest-like combinatorial optimization among the top 6 candidate set (Section 2.2) is used to guard against a circular comparison. The specificity claim is validated against an independent expert curated resource (Section 3.4).

## 2 Materials and Methods

### 2.1 Datasets

The analysis was conducted using three publicly available annotated atlases. The Human Lung Cell Atlas Core (HLCA; Sikkema et al., 2023) is composed of 584,944 cells, constituting 61 unique cell type clusters at the *ann_finest_level* annotation. The HLCA Core was generated using 10x Genomics droplet-based single-cell RNA sequencing. Cluster size ranges from 28 to 68,487 cells (2,446-fold range) (https://cellxgene.cziscience.com/collections/6f6d381a-7701-4781-935c-db10d30de293). The adult human kidney atlas (Lake et al., 2023) contains 107,701 single-cell and single-nucleus transcriptomes forming 76 clusters at the *subclass*.*l3* level. These data were also derived using the 10x Genomics droplet-based RNA-seq platform, and cluster sizes range from 6 to 8,551 cells (1,425-fold range) (https://cellxgene.cziscience.com/collections/bcb61471-2a44-4d00-a0af-ff085512674c). The final dataset used was the human middle temporal gyrus dataset (Brain MTG; Hodge et al., 2019). The Brain MTG dataset contains 15,603 nuclei, encompassing 75 clusters, generated by plate-based Smart-seq v4. Cluster sizes range from 9 to 2,284 cells (254-fold range) (https://brain-map.org/our-research/cell-types-taxonomies/cell-types-database-rna-seq-data). Preprocessing and label handling are described in Supplementary Materials S1.1.

### 2.2 Marker selection

DE markers were identified through Scanpy’s *rank_genes_groups* using *method = ‘wilcoxon’, reference = ‘rest’*, and *tie_correct = False* (Scanpy’s defaults and recommendations). NS-Forest markers were generated using NS-Forest v4.1 (https://github.com/NLM-DIR/NSForest) using default parameters. Full parameters for both methods are given in Supplementary Materials S1.2.

Two DE marker sets were defined per cluster to test Wilcoxon’s realistic and best-case performance. DE-practiced (DEprac) takes the top-k positively enriched genes by Wilcoxon score, where k is the number of markers NS-Forest reported for that cluster. This is DE in its typical application when using Seurat’s or Scanpy’s default and recommended marker selection settings. DE-optimized (DEopt) takes the top six positively scored DE genes as a candidate pool and submits them to NS-Forest’s combination search that maximizes classification F-beta (Supplementary Materials S1.2).

DEopt grants DE the same combinatorial optimization advantage that NS-Forest provides and represents a DE ideal marker set rather than its typical use. Both DE marker sets and the NS-Forest set were scored identically by the evaluator described in Section 2.4.

### 2.3 Effect-size ranking (DEauc)

The test statistic returned by *rank_genes_groups* for *method = ‘wilcoxon’* is

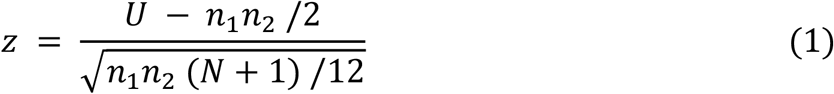

where *N* is the total number of cells, *n*_1_ is the target cluster size, and *n*_2_ is the background comparison group (*n*_2_ = *N* - *n*_1_). *U* is the Mann-Whitney statistic, which quantifies the number of target-versus-background cell pairs in which the target cell has a higher gene expression, with tied pairs counted as one half. The denominator is the standard deviation of *U* under the null hypothesis. Therefore, at a fixed underlying effect, |*z*| grows with 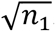. Inverting the equation recovers the rank-based effect size to analyze the win-probability independent of the size contamination (DEauc).

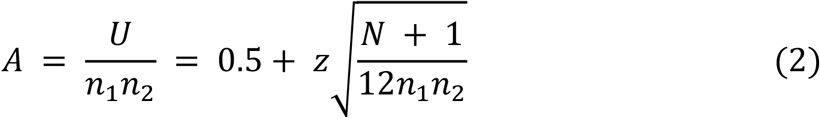

*A* is the probability that a randomly drawn cell from the target cluster expresses a gene more highly than a randomly drawn cell from the rest of the atlas. *A* is a discrimination proportion bounded on [0,1] and equals 0.5 for a non-discriminating gene. This discrimination proportion is numerically identical to the area under the receiver operating characteristic (AUROC) curve - a known property in statistics (Hanley and McNeil, 1982). Unlike *z*, it does not grow with sample size at a fixed effect. DE candidates ranked by *A* rather than *z* will be referred to as DEauc.

Because *n*_1_, *n*_2_, and *N* are fixed within a cluster, *A* is an increasing transformation of *z*. Ranking genes by *A* yields an identical ordering, and therefore an identical marker set as ranking by *z* (Supplementary Materials S1.3). Because of this, DEauc nominates the same genes as DEprac, meaning its recall, precision, and on-target fraction are identical. Effect-size ranking removes the size dependence in how genes are scored without altering which genes are selected.

### 2.4 Evaluation metrics

All three marker sets (DEprac, DEopt, and NS-Forest) were scored for cell type classification performance using standard measures of F-beta, precision, and recall using the classification evaluator built into NS-Forest. Binary classifiers for each marker were built using depth-1 decision tree split (Gini criterion, high side positive). Recall, precision, and F-beta (*β* = 0.5) were computed by the same evaluator for all marker sets. F-beta is a weighted F1 score, where *β* = 0.5 weights precision twice as important as recall, mitigating inflated false negatives due to the dropout artifact in scRNA-seq experiments.

**On-target fraction (OTF)** is the primary metric used to quantify specificity. For a gene *g* and target cluster *T*, OTF is the median expression, *m*, of *g* in *T* divided by the summation of its per-cluster medians across all clusters.

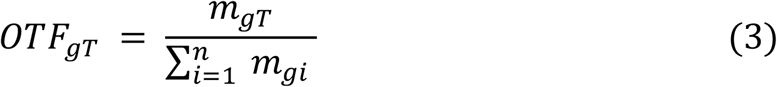

Because OTF is a specificity metric built from medians rather than rank sums, OTF carries no structural size term and is a post-hoc evaluator for both methods. Its denominator sums over all clusters, so cluster OTF values are comparable within an organ, but not across organs with different cluster counts.

OTF differs from the classification metrics above in how it weights the atlas. Precision and recall are cell-weighted, as both are computed from counts of individual cells, while OTF is cluster-weighted due to the nature of its denominator. Each cluster contributes one median per cluster regardless of the cluster’s size, meaning a gene expressed throughout a lineage of small, related clusters is penalized heavily by OTF but only slightly by precision. This allows OTF to capture the cluster-wide specificity and is in theory robust to cluster sizes.

### 2.5 Downsampling and simulation

For the downsampling experiment, cells were progressively capped per cluster at a ladder of values ranging from uncapped down to 50 cells for lung and 20 cells for kidney and brain. DE was recomputed at each rung across 5 to 20 random seeds per cap; more seeds were utilized at smaller caps where between run variation is largest. Cluster identity was held constant while *N* was varied.

To test whether cluster size alone is responsible for DE score inflation, we built a simulated dataset in which marker quality is fixed. We generated 42 synthetic clusters, 14 log-spaced sizes from 30 to 30,000 cells with one cluster per marker quality tier at each of the 14 sizes. Gene expression was drawn from a negative binomial distribution (dispersion *θ* = 2). We defined three tiers of marker-quality defined by off-target expression: high (off-target mean = 0.05), medium (0.95), and low (1.2) (Supplementary Figure S1). Each cluster was assigned exactly one marker of a single quality tier, so marker quality and cluster size vary independently across the grid. All planted markers had a mean in-target expression of 6.0 regardless of their quality, followed by 20% dropout. The realized on-target fractions of each quality tier were 1.00, 0.53, and 0.14 respectively. This was simulated over 15 random seeds.

### 2.6 Statistical analysis

Pearson correlations (*r*) were computed against *log*_10_ cluster sizes with Fisher *z* 95% confidence intervals for each organ. Spearman rank correlations (*ρ*) are also reported. Partial correlations were computed to determine if the DE-size relationship is mediated by classification performance. Paired method comparisons used Wilcoxon signed-rank test using rank-biserial correlations as the effect size. All differences below 0.01 were counted as ties. Note that the Wilcoxon signed-rank test used for paired comparisons is distinct from the Wilcoxon ranked-sum test that is being evaluated as a marker selection method. Confidence intervals on partial correlations and median metrics were calculated using bootstrap resampling of clusters (2,000 replicates). Analyses used Python 3.12 with Scanpy 1.12.1, NumPy, pandas, and SciPy. NS-Forest v4.1 was installed from source (https://github.com/NLM-DIR/NSForest). Simulation and analysis scripts are available at https://github.com/NLM-DIR/nsforest-vs-de-marker-selection.

## 3 Results

### 3.1 Wilcoxon DE scores scale with cluster size across organs

Cluster sizes in all three atlases (lung, kidney, brain) span two to three orders of magnitude with medians more than an order of magnitude smaller than the largest cluster (Figure 1A).

**Figure 1.**
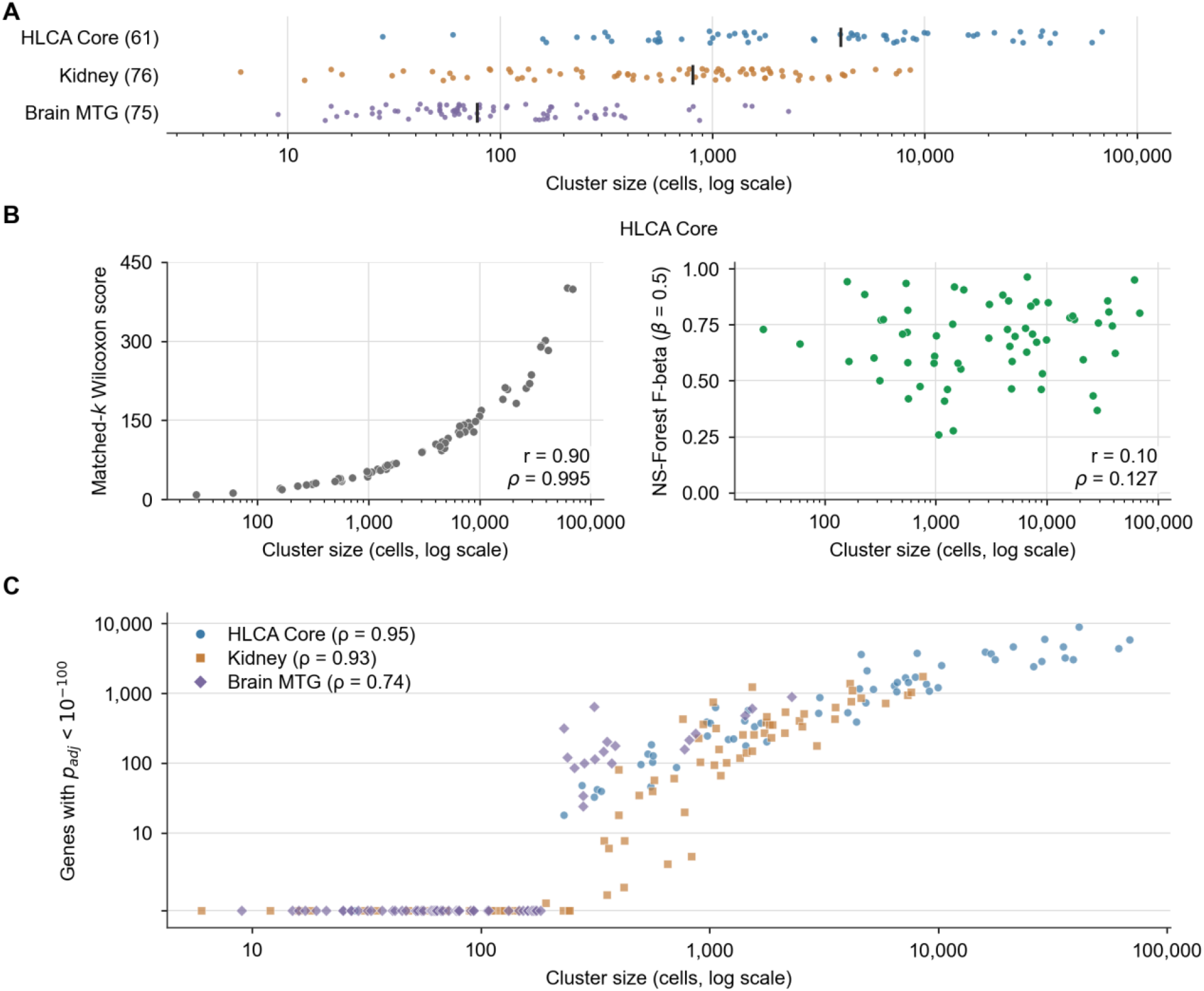
Wilcoxon rank-sum DE scores scale with cluster sizes across all three datasets while NS-Forest F-beta does not. **(A)** Distribution of cluster sizes (number of cells, log10 transformed) in each atlas: HLCA Core (lung), 61 clusters spanning 28 - 68,487 cells (2,446-fold range); Kidney, 76 clusters spanning 6 - 8,551 cells (1,425-fold range), Brain MTG, 75 clusters spanning 9 - 2,284 cells (254-fold range). Vertical ticks mark median cluster sizes in each atlas. **(B)** HLCA Core, one point per cluster. Left: matched-k mean Wilcoxon score vs. log cluster size (k set per cluster by NS-Forest’s final marker count): *r* = 0.903 (*ρ* = 0.995), Right: NS-Forest F-beta (*β* = 0.5) vs. log cluster size on the same clusters: *r* = 0.10 (*ρ* = 0.127) (Kidney and Brain MTG in Supplementary Figure S2). Y-axes bounded to [0,1] by construction of F-beta. **(C)** Number of genes with adjusted *p*-value < 10^−100^ per cluster versus cluster size in all three datasets, axes log-scaled. Unrankable gene counts rise with cluster size across all atlases (*ρ* = 0.95, 0.93, 0.74). Clusters with no unrankable genes fall to the axis floor.

Under these conditions, the matched-k (same number of k genes in DE and NS-Forest) mean Wilcoxon score scaled with log cluster size in every organ (Figure 1B, left and Supplementary Figure S2). The near perfect positive Spearman values indicate that the ordering of clusters by DE score is nearly identical to the order of clusters by size.

NS-Forest’s F-beta was computed on the same clusters and showed virtually no dependence on cluster size (Figure 1B, right and Supplementary Figure S2).

This effect is not unique to Wilcoxon. Across the lung, kidney and brain datasets, three additional differential expression tests were performed (Supplementary Table S1). All four tests showed substantial scaling with cluster size. The most widely used test, Wilcoxon, is the one whose rankings are most tightly correlated with cluster size.

Ranking by adjusted *p*-values in large clusters fails for a related but separate reason. The statistical power of a one-versus-rest comparison in scRNA-seq data grows with cluster size and, as a result, many genes in large clusters are driven to adjusted *p*-values so small that the differences between them carry no meaningful information. Genes with an adjusted *p*-value below 10^−100^ cannot accurately be ordered against each other, and *p*-value saturation appears in clusters from 192 cells upward (Figure 1C). Therefore, ranking genes by *p*-value deteriorates into arbitrary tie-breaking for clusters with the most data. Because the ordering among saturated genes no longer reflects any differences in effect, the residual separation between their ranks is due to sampling variation, meaning gene rankings are least stable in the clusters with the most data.

*P*-value is monotone decreasing in |*z*|, meaning it inherits the size dependence and adds to the saturation effect.

### 3.2 The dependence is a property of the statistic and not of marker quality

Four separate lines of evidence establish the DE-size relationship as a property of the underlying statistical methodology rather than a reflection of genuinely better markers in large clusters.

#### Controlling for marker quality leaves the DE-size relationship intact

If larger clusters had more distinct markers, then controlling for classification performance through NS-Forest’s F-beta should weaken the correlation. Partial correlation tests this by asking how much of the DE-size correlation remains after F-beta’s contribution to each has been accounted for (Figure 2A and Supplementary Figure S3). F-beta is a suitable control variable for this test because it is composed of proportions that exclude true negatives, so it does not scale with sample size as the DE score does. Precision does rise modestly with cluster prevalence, making F-beta weakly size-dependent. With this, controlling for F-beta removes a small amount of size-related variance, therefore making this test strict. When removing the effect of F-beta, *r*(DE, size) remained virtually unchanged in all organs (lung, 0.90 → 0.91; kidney and brain in Supplementary Figure S3), suggesting high correlation between DE and cluster size remained regardless of classification performance. A reciprocal partial correlation was computed between DE score and F-beta, controlling for size, resulting in a weak correlation across all organs (lung 0.21 → 0.29; kidney and brain in Supplementary Figure S3). Together, these results indicate that the DE-size relationship is not mediated by classification performance.

**Figure 2.**
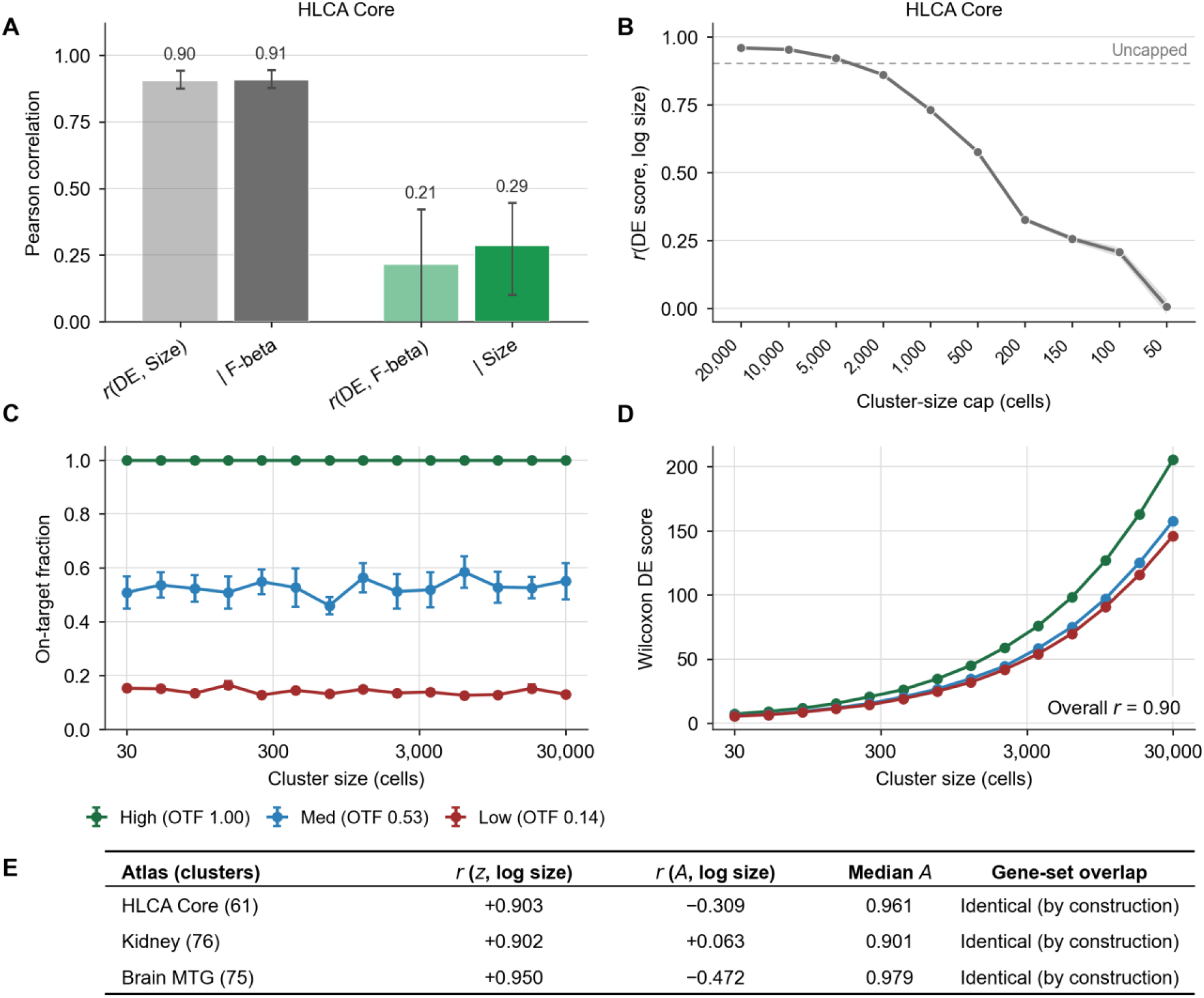
The cluster size dependence is a property of the Wilcoxon rank-sum statistic and is not due to better marker selection in large clusters. **(A)** Pearson correlations of HLCA Core with bootstrap 95% confidence intervals. Bars represent controlling for variable in partial correlation. NS-Forest’s F-beta was removed by partial correlation, leaving *r*(DE, size) essentially unchanged (0.90 → 0.91). A reciprocal partial correlation was conducted showing that the relationship between DE and F-beta is weak with or without controlling for size (0.21 → 0.29) (Kidney and Brain MTG Supplementary Figure S3). **(B)** Progressively capping cells per cluster in HLCA Core holds cell identity and profile fixed while cluster size is reduced. *r*(DE, size) declines monotonically across the cap range. Dashed line represents uncapped correlation. Kidney and brain MTG in Supplementary Figure S3. **(C**,**D)** Simulation with marker quality fixed (42 clusters: three off-target expression tiers x 14 log-spaced clusters ranging 30 - 30,000, run over 15 seeds). **(C)** On-target fraction vs log cluster size. Tiers shown in green (high), blue (medium), and red (low). **(D)** Wilcoxon score vs log cluster size (overall *r* = 0.90). **(E)** Table shows z, the Wilcoxon rank-sum standardized statistic, is correlated with log cluster size. A = U/(n1n2) is a rank-based effect size transformation of z bounded to [0,1] with 0.5 indicating no discrimination. Correlations are Pearson *r* against log10 cluster size, computed on the same matched-k mean per cluster as prior analyses.

#### Equalizing cluster sizes removes DE-size correlation

Progressively capping cells per cluster holds cell identity constant while only varying cluster size *n*. The observed correlation between matched-k DE score and log cluster size decreased monotonically across the cap range in every organ (Figure 2B and Supplementary Figure S3). The endpoint correlations of this ladder order were *r* ≈ 0.01 (lung), 0.26 (brain), and 0.35 (kidney), which is expected if the statistic is tracking cluster size. Some residual variation remained and is explained in Supplementary Materials S1.4.

#### Correlation reproduces when specificity is fixed by construction

We built a simulation of synthetic scRNA-seq data where marker quality is defined into three tiers by the generative model through off-target expression levels and does not vary with cluster size (14 cluster sizes, log spaced 30 to 30,000). OTF correctly distinguishes the three marker quality tiers and remains flat across the range of cluster sizes, confirming it recovers the designed ordering (Figure 2C and Supplementary Figure S1). Wilcoxon scores climb as cluster size grows regardless of the marker quality (Figure 2D). A marker that is of low quality and expressed widely off-target in a large cluster outscores an on-target marker of high quality in a small cluster by the Wilcoxon score.

#### Effect-size ranking eliminates the dependence when removing the sample-size term

*A* is the proportion of target-background cell pairs (*n*_1_*n*_2_) in which the target cell has a higher gene expression value than an off-target cell (*U*) (Equation 2). This is a ratio fixed by how often the gene’s expression rank is higher in the target cluster than the background. The statistic *z* uses this quantity and rescales it by the null variability, which grows more slowly than the pair count. At a fixed, underlying effect, *z* grows with 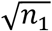, while *A* does not.

Ranking clusters by *A* rather than *z* removes the positive size dependence, yet returns an identical set of genes, verified in every cluster across all organs (Figure 2E). The size dependence is therefore attributable entirely to the sample-size term in the denominator (Equation 1), and not to any property of the genes themselves.

### 3.3 DE nominates genes that are less specific

The scaling artifact is controllable, but the genes it nominates are not.

Across the four performance metrics (Figure 3A), the methods clearly separate on one: OTF. NS-Forest has the highest median OTF of 0.580, 0.687, and 0.364. DEprac, the marker set that any standard workflow using Seurat’s or Scanpy’s default and recommended settings produces, performed the worst in OTF with medians of 0.204, 0.166, and 0.083. DEopt, the enhanced version of DE, did not recover OTF. The three cell-weighted metrics did not separate the methods consistently. F-beta favored NS-Forest in lung (*p* = 0.018), tied in kidney, and favored DEopt in brain (*p* = 0.020). Median precision was lower for DEprac than the other two combinatorial methods, while the median recall was highest for DEprac in all three organs. This inversion is expected, as without set optimization, DEprac favors broadly expressed genes at the cost of precision.

**Figure 3.**
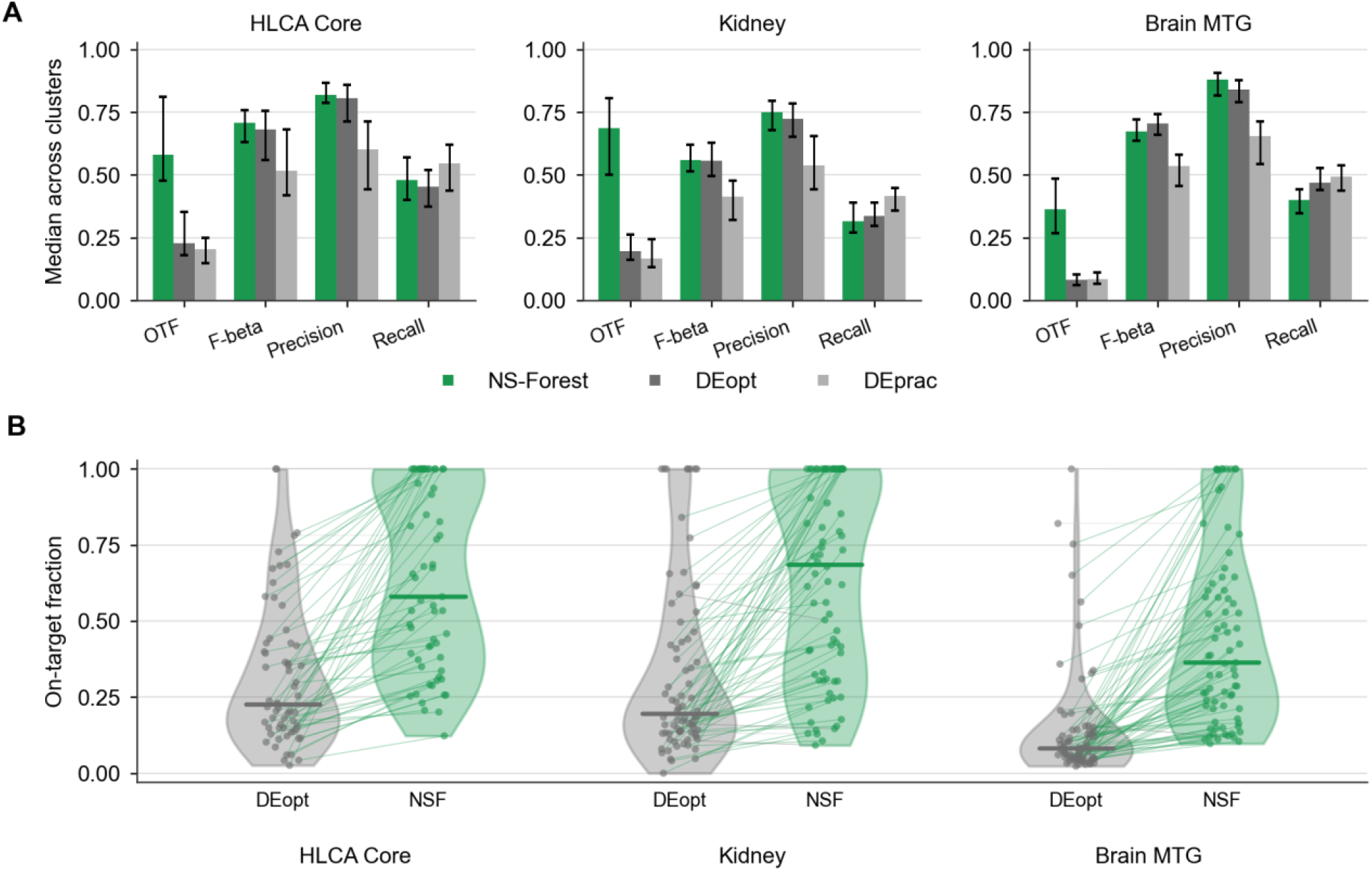
Markers nominated by DE are substantially less on-target than NS-Forest nominated markers, unrecoverable by running top DE genes through combinatorial optimization. **(A)** Median across clusters of four metrics comparing NS-Forest, DEopt, and DEprac (default DE) with bootstrap 95% confidence intervals. **(B)** Per-cluster on-target fraction for optimized DE (DEopt) vs. NS-Forest, one point per cluster, median marked with horizontal tick for each method. Medians (NS-Forest vs DEopt): HLCA Core 0.580 vs 0.227; kidney 0.687 vs 0.195; brain MTG 0.364 vs 0.082.

On-target fraction, which measures how concentrated a gene’s expression is within its target cluster, contrasted considerably between methods across all organs (Figure 3B). NS-Forest markers showed higher on-target specificity in 54 of 61, 60 of 76, and 71 of 75 clusters against DEopt (paired signed-rank, all *p* < 10^−10^, rank-biserial 1.00, 0.98, 0.99). These differences indicated that more specific marker sets could be obtained than the DE nominated genes.

The methods remain close on F-beta and precision due to these metrics being cell-weighted, meaning off-target expression in small, related clusters contributes too few cells to substantially move them.On-target fraction is cluster-weighted, so that same expression enters the denominator in full and causes the clear separation between methods. Optimization improved DE’s classification performance through precision and F-beta, but left OTF essentially unchanged relative to the naive version of DE. Both rescaling the statistic and optimizing for classification did not change DE’s marker candidate selection to improve specificity.

### 3.4 DE picks class markers that are highly abundant

To illustrate the implication of DE’s non-specificity, our results from the largest atlas (HLCA) were checked against an independent, expert curated resource: LungMAP cell cards (Sun et al., 2022). LungMAP is treated as a validator for markers found by these methods, but not used as a specificity benchmark as it does not report specificity metrics.

For arterial endothelial cells (EC), NS-Forest selected *DKK2* (OTF 1.00), which LungMAP lists among its known arterial endothelial markers. DE selected *CLDN5* (OTF 0.149), a marker in which literature describes as pan-endothelial, is not listed in LungMAP as arterial specific (Figure 4A).Further analysis showed that *CLDN5* was highly expressed in a considerable fraction of cells within all eight endothelial clusters in the atlas (Figure 4B).

**Figure 4.**
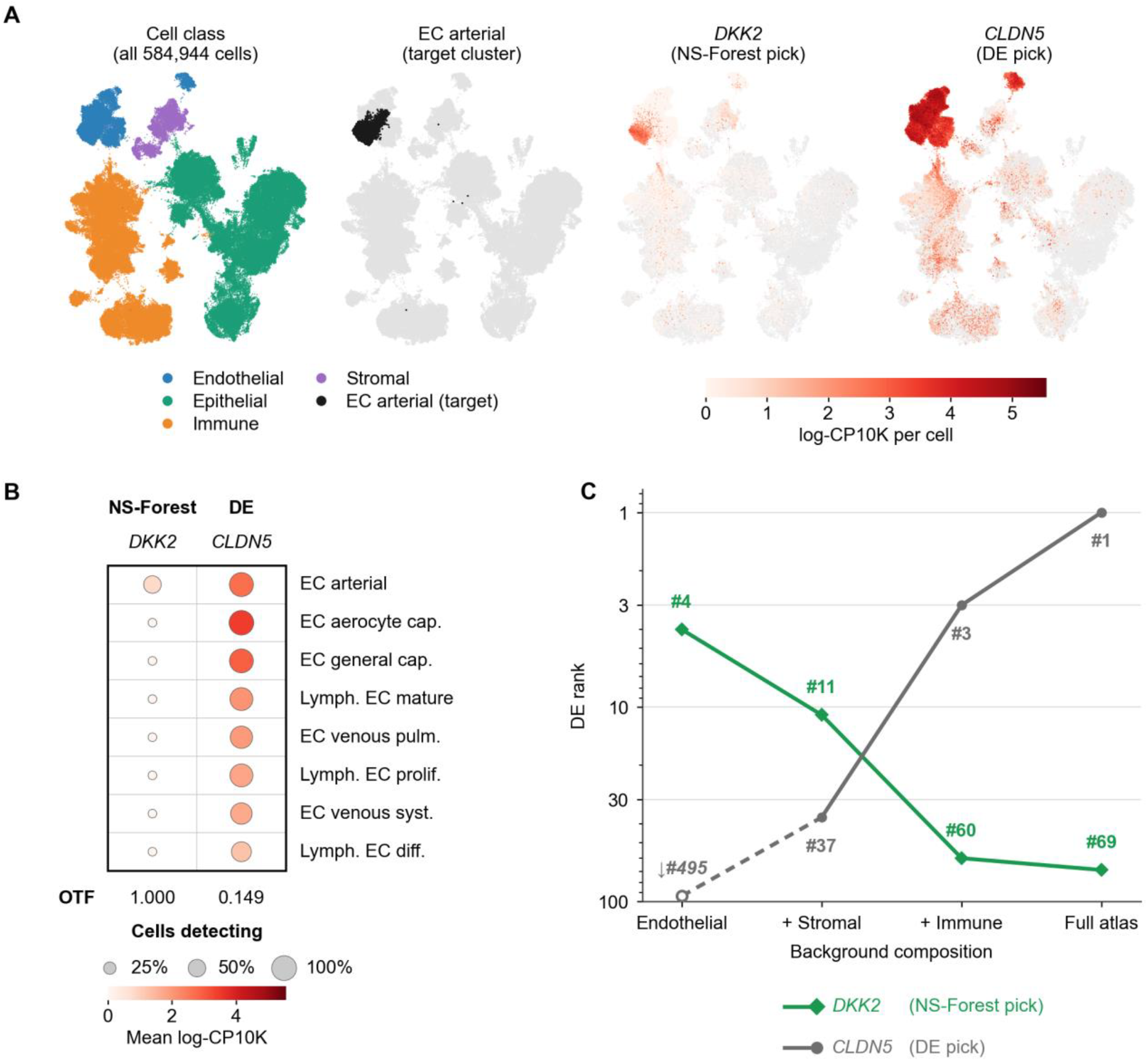
The DE-nominated marker for EC arterial cells is widely expressed across the endothelial lineage, and its ranking is dependent on its background comparison group. **(A)** HLCA Core UMAP. Left: Class structure of HLCA Core. Middle left: EC arterial target cluster. Middle right: per-cell expression of *DKK2*, the NS-Forest marker selected. Right: per-cell expression of *CLDN5*, DE’s top ranked marker using a shared color scale. **(B)** Expression of *DKK2* and *CLDN5* across the eight endothelial clusters in HLCA Core. Area of each dot represents the fraction of cells expressing the gene and color represents the mean log-CP10K on a shared absolute scale with no rescaling. *CLDN5* is expressed higher in an off-target cluster (EC aerocyte capillary cells, 3.456) than its own target cluster (EC arterial cells, 2.700). On-target fraction was computed against all 61 clusters. **(C)** DE rank trajectory of EC arterial selected markers as the background group widens across the atlas hierarchy (endothelium only, adding stroma, adding immune, full atlas). The two genes invert between rungs 2 and 3 where the background stops being predominantly endothelial.

We followed the DE rank of *CLDN5* and *DKK2* as we widened the background group, beginning with the endothelial class alone, then adding one cell type class at a time until all four classes were in the background (Figure 4C). Against endothelial clusters alone, DE ranked *CLDN5* 495th. When expanded to the full atlas *CLDN5* ranked 1st. *DKK2* moves in the opposite direction, from 4th to 69th. Neither gene expression nor target cluster changed between comparisons, only the composition of the comparison group changed.

A similar pattern is observed in submucosal gland serous cells. NS-Forest selected *STATH* (OTF 1.00) and DE selected *SLPI* (OTF 0.09), neither of which appears in LungMAP’s marker list (Supplementary Figure S4). NS-Forest selects the marker that is much more specific to the target cluster, while again DE selects a gene that is broadly expressed across the entire epithelial class.

Finally, in AT2 cells, the second largest cluster in the entire dataset, NS-Forest selected *SFTPA1* with DE selecting *SFTPB* and LungMAP listing *SFTPC* (Supplementary Figure S5). All three of these genes work in conjunction as pulmonary surfactant genes, yet both NS-Forest and DE pick the more specific marker than listed in LungMAP. This is why LungMAP is treated as a validator of marker identity and not a benchmark of specificity. In all three cases, NS-Forest picked a more on-target marker gene than DE.

## 4 Discussion

Two clear drawbacks to DE-based marker gene identification were identified through this analysis, with two distinct consequences. The first is a structural issue in the Wilcoxon rank-sum test.

Wilcoxon’s test statistic (*z*) scales with target cluster size, making scores incomparable across clusters. Second, the genes DE proposes systematically fail to capture specificity within the target cluster and tend to be expressed in closely related cell types within a cell lineage. Panels that are assembled using ranking by DE scores inherit a bias toward abundant cell types, and their markers leak into related clusters.

The first drawback can be controlled through reranking by a proportion, *A*. However, this does not improve specificity as *A* is a monotone transformation of *z* within each cluster. DEprac, the marker genes produced by Seurat’s and Scanpy’s default and recommended workflows, was the least specific of all three marker sets. DEopt failed to improve in specificity even when granted the same combinatorial optimization NS-Forest uses. This suggests that specificity is a property of the genes that enter the candidate pool, and neither rescaling the ranking nor optimizing for classification performance can improve the candidate pool obtained using Wilcoxon.

A single property of classification explains several of the observations from this research at once. In one-versus-rest testing, such as Wilcoxon’s, target cluster size determines how rare the positive class is, ranging here from a few cells to tens of thousands. Recall (TP/(TP+FN)) is computed within target cells alone, meaning the size of off-target clusters will not affect this metric. The same is not true for precision (TP/(TP+FP)), as false positives are drawn from the rest of the atlas, while true positives will scale with cluster size. F-beta, a weighted harmonic mean of precision and recall, is an effective measure of a marker’s classification power because true negative (TN) counts are not in its equation. TNs are the number of off-target cells in an atlas, which is often a very large number (even much larger than the largest cluster), and would dominate any measure if included. For this reason, DE-based measures (*z* or *A*), which consider all cells regardless of true/false positives/negatives, are not effective for selecting markers for cell classification. Classification metrics, however, are not the only way to evaluate a marker. Precision and F-beta ask whether a marker gene can identify its target cluster within an atlas of heterogeneous cell types. On-target fraction measures whether a gene’s expression is concentrated in a cell type rather than distributed across the atlas and is therefore a complementary measure of specificity to precision and F-beta.

Wilcoxon is being used to do a job it was never designed to do. Within a single cluster, its ranking is accurate to its equation, which is why ranking by *A* returns the same top genes. What it cannot support is comparing its scores across clusters of different sizes, which is precisely what marker gene selection requires.

### 4.1 Limitations

We examined three organs, spanning two capture technologies. We took the annotated labels from each of the authors as ground truth, so annotation errors populate both methods equally. On-target fraction originates from the comparator method, NS-Forest, which is why we examined the central claim against the external database of LungMAP, which was only available for one of the three organs. We did not vary DE filters to apply minimum-detection or fold-change thresholds before ranking, so the conclusions are based on the unfiltered, default products of DE rather than a variant workflow. The discussion in this study only applies when using DE for cell-type marker discovery and does not apply when using DE analysis to compare samples between conditions (e.g., healthy versus diseased).

### 4.2 Recommendations

We suggest the following recommendations when it comes to current practices for identifying marker genes for scRNA-seq atlases. First, do not rank marker gene candidates across clusters of unequal size by a standardized test statistic or adjusted *p*-value. Every DE test we examined scaled with cluster size, with the field’s default method of Wilcoxon scaling the steepest. Adjusted *p*-values fail as they saturate at an incomparable floor in the clusters with the most data. Second, rescaling the statistic is not a solution. Ranking based on effect size *A* = *U*/(*n*_1_*n*_2_) removes the size dependence but returns the same list of candidate genes. No adjustment to the test statistic will recover specificity. Third, it is important to report a proportion-based specificity metric alongside any marker panel to quantify the marker gene’s expression within-target. Last, where the goal is to identify cell types within a population, selection methods built on classification are preferred. When motivated by identification rather than detection, trading off recall for improved precision is appropriate. The diagnostic here is computationally inexpensive and can be run on any annotated atlas.

## 5 Data availability statement

The datasets used for this analysis are publicly available. The Human Lung Cell Atlas Core (Sikkema et al., 2023) is available at: https://cellxgene.cziscience.com/collections/6f6d381a-7701-4781-935c-db10d30de293. The adult human kidney atlas (Lake et al., 2023) is available at: https://cellxgene.cziscience.com/collections/bcb61471-2a44-4d00-a0af-ff085512674c. The human middle temporal gyrus dataset (Hodge et al., 2019) is available at: https://brain-map.org/our-research/cell-types-taxonomies/cell-types-database-rna-seq-data. All analysis code, including marker selection, evaluation, simulation, and figure construction is available at https://github.com/NLM-DIR/nsforest-vs-de-marker-selection.

## 6 Ethics statement

Ethics approval was not required for this study as all data was collected through previously published and publicly available de-identified human single-cell RNA sequencing datasets (Hodge et al., 2019; Lake et al., 2023; Sikkema et al., 2023). No new data was generated. Ethical approval was obtained by the original source studies.

## 7 Author contributions

KD: Conceptualization, Data curation, Formal analysis, Investigation, Methodology, Software, Validation, Visualization, Writing – original draft, Writing – review & editing. DP: Investigation. RHS: Conceptualization, Funding acquisition, Supervision, Validation, Writing – review & editing. YZ: Conceptualization, Data curation, Funding acquisition, Investigation, Project administration, Resources, Software, Supervision, Writing – review & editing.

## 8 Funding

This research was supported by the Intramural Research Program of the National Institutes of Health (NIH). The contributions of the NIH authors are considered Works of the United States Government. The findings and conclusions presented in this paper are those of the authors and do not necessarily reflect the views of the NIH or the U.S. Department of Health and Human Services.

## 9 Conflict of interest

The authors declare that the research was conducted in the absence of any commercial or financial relationships that could be construed as potential conflicts of interest.

## 10 Generative AI statement

The authors declared that generative AI was used in the creation of this manuscript. Generative AI was used during the editing process of the text. The original and final draft was written by the first author and edited by all authors. Generative AI was used to improve the clarity and grammar of the final draft. The author reviewed and edited all the AI prompted text and takes full responsibility for the content of the publication.

## Notes

https://github.com/NLM-DIR/nsforest-vs-de-marker-selection

### Competing Interest Statement

The authors have declared no competing interest.

