## Supplementary Material for "Limitations of differential expression for cell-type marker discovery in single-cell RNA sequencing atlases"

### S1. Supplementary Data

#### S1.1 Data preprocessing

Gene expression matrices were used as log-normalized counts-per-10,000 (log-CP10K). No clustering was recomputed and published cell type cluster labels were accepted throughout the analysis. No cells or nuclei were reassigned from their annotated cluster. The distribution of cluster size across the three datasets is shown in Figure 1A.

#### S1.2 Marker selection parameters

DE markers were identified through Scanpy's *rank_genes_groups* using *method = 'wilcoxon'*, *reference = 'rest'*, and *tie_correct = False* (Scanpy's defaults). These conditions yielded a one-versus-rest comparison for each target cluster against all remaining cells in the atlas. Since positive markers are more useful in downstream research, only positively scored DE genes were considered.

NS-Forest markers were generated using NS-Forest v4.1 (Liu et al., 2024) using default parameters. A marker set for each cluster was computed on the same annotation column and the same log-normalized expression matrix used for DE. NS-Forest only reports positive markers.

In the combinatory stage of the candidate genes that NS-Forest and DEopt undergo, the top 6 candidates undergo a combination search to optimize F-beta in its final set. This equates to 6!=720 total combinations searched. This cap of six candidates is set to maximize combination searches while preserving computing.

The returned sets generally contain one to four genes, and averages out to 2.1 genes per set. The reduced number of genes in the final set is a byproduct of $\beta = 0.5$, as each additional gene added to the set can drop true positives along with false positives, and therefore raising precision but decreasing recall, and F-beta weights both. Set size is therefore selected on the maximum F-beta combination and set size varies by cluster.

#### S1.3 Effect size recovery

The rank-based effect size $A$ was recovered from Scanpy's returned z statistic by inverting Equation 1. The recovery is precise under Scanpy's default of no tie correction, and all recovered $A$ values fell within [0,1] in every organ. The recovery was verified when $U$ was computed manually, agreeing to within $7.3 \times 10^{-9}$.

#### S1.4 Residual variation

The downsampling ladder performed in Figure 2B and Supplementary Figure S3 progressively caps the maximum number of cells contributed to each cluster at a fixed ceiling, drawing without replacement for clusters exceeding the cap and leaving clusters under the cap unchanged.

Because capping truncates the upper tail rather than equalizing sizes, the smallest cap per atlas is set by its smallest cluster, and some residual size-variation remains: 1.8-fold in lung (smallest cluster of

28 with all other clusters set to a max-cap of 50), 2.2-fold in brain (9 with a max-cap of 20), and 3.3-fold in kidney (6 with a max-cap of 20).

Residual correlation orders with residual size variation across all three atlases. Lung had the smallest floor-to-cap ratio and retained nearly no correlation between DE score and log cluster size, while kidney had the largest floor-to-cap ratio and retained the most correlation.

### Supplementary Figures

#### Supplementary Figure S1

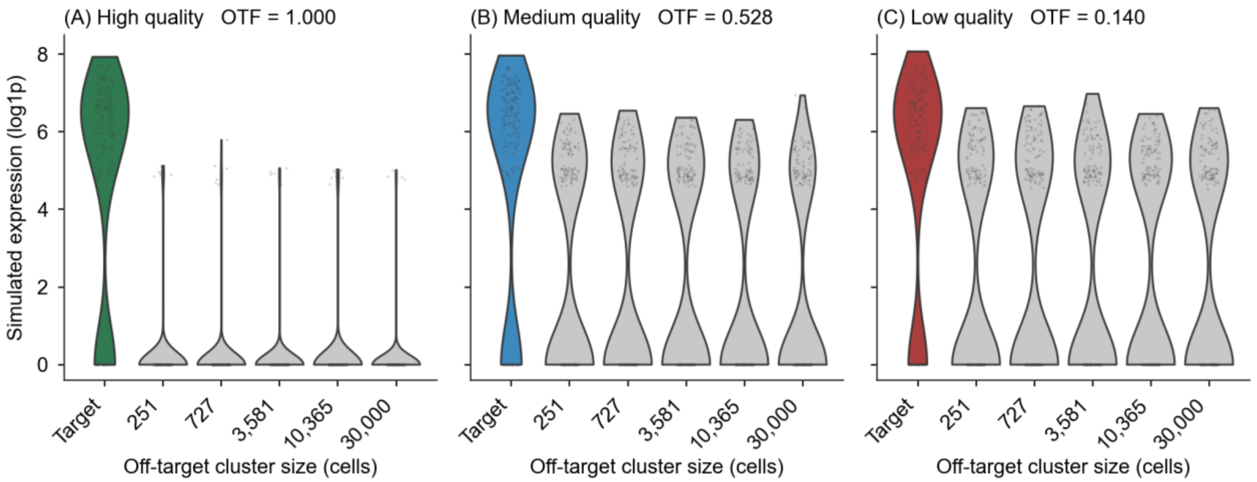


**Supplementary Figure S1. Simulation marker quality expression profile across specificity tiers.** Each panel shows the expression profile for the planted marker in one simulated cluster for each of the specificity tiers. Violin plots are density estimates with points overlaid representing individual cells. All violins are subsampled to 200 for easy visualizations while off target clusters are labeled by their full, true size. Tiers are differentiated by the markers off-target expression, with medium and low quality markers expressed highly in off-target clusters. **(A)** High quality marker: off-target mean expression 0.05. **(B)** Medium quality marker: off-target mean expression 0.95. **(C)** Low quality marker: off-target mean expression 1.20. Marker expression in the target cluster is identical across tiers by construction for comparison (in-target mean expression = 6.0, 20% dropout, negative binomial dispersion $\theta = 2$). Realized on-target fraction shown in each tier of marker quality.

**Supplementary Figure S2**

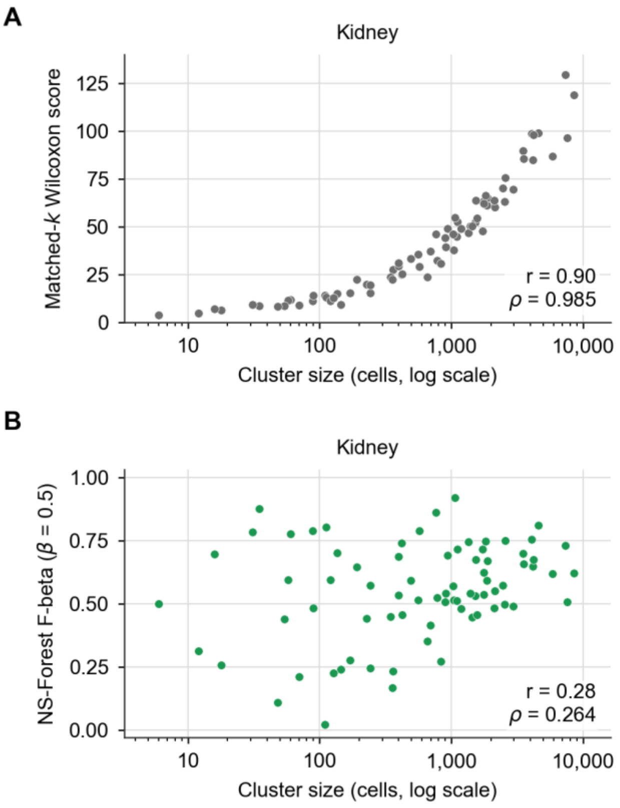


**Supplementary Figure S2. The DE-size relationship replicates in kidney and brain datasets while NS-Forest's remains independent.** Per-cluster scatter for the two atlases not shown in Figure 1B. **(A)** Matched-k mean Wilcoxon score vs. log cluster size, one point per cluster (k set per cluster by NS-Forest's final marker count): kidney (76 clusters) $r = 0.902$ ($\rho = 0.985$), Brain MTG (75 clusters) $r = 0.950$ ($\rho = 0.994$). Y-axis is free because of the unbounded nature of the Wilcoxon statistic. **(B)** NS-Forest F-beta ($\beta = 0.5$) vs. log cluster size on the same clusters: kidney $r = 0.28$ ($\rho = 0.264$); Brain MTG $r = 0.21$ ($\rho = 0.209$). Y-axes are fixed to [0,1] by construction of F-beta. Pattern matches observed results for HLCA Core in Figure 1B.

**Supplementary Figure S3**

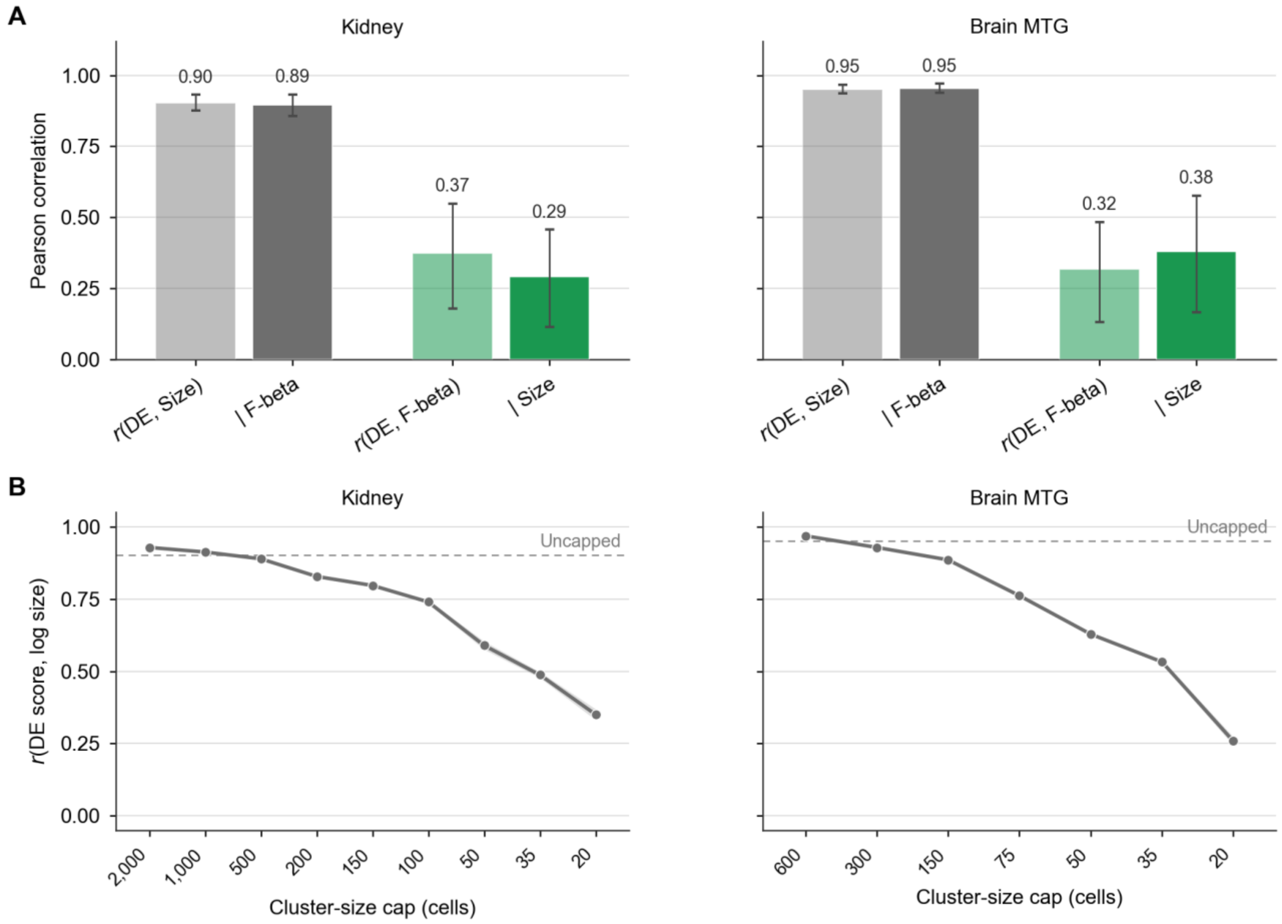


**Supplementary Figure S3. Cluster size dependence is matched in kidney and brain atlases. (A)** Pearson correlations with bootstrap 95% confidence intervals. NS-Forest F-beta was removed by partial correlation, leaving *r*(DE score, log cluster size) essentially unchanged in both atlases (kidney 0.90 → 0.89, brain MTG 0.95 → 0.95). A reciprocal partial correlation was conducted on the two datasets showing that the relationship between DE and F-beta is weak with or without controlling for size (kidney 0.37 → 0.29, brain MTG 0.32 → 0.38). Matches results seen in HLCA Core. **(B)** Progressively capping large clusters in kidney and brain holds cell identity constant and expression profile fixed while varying cluster size. *r*(DE score, log size) declines monotonically as the cap tightens in both atlases. Dashed lines represent the uncapped correlation.

**Supplementary Figure S4**

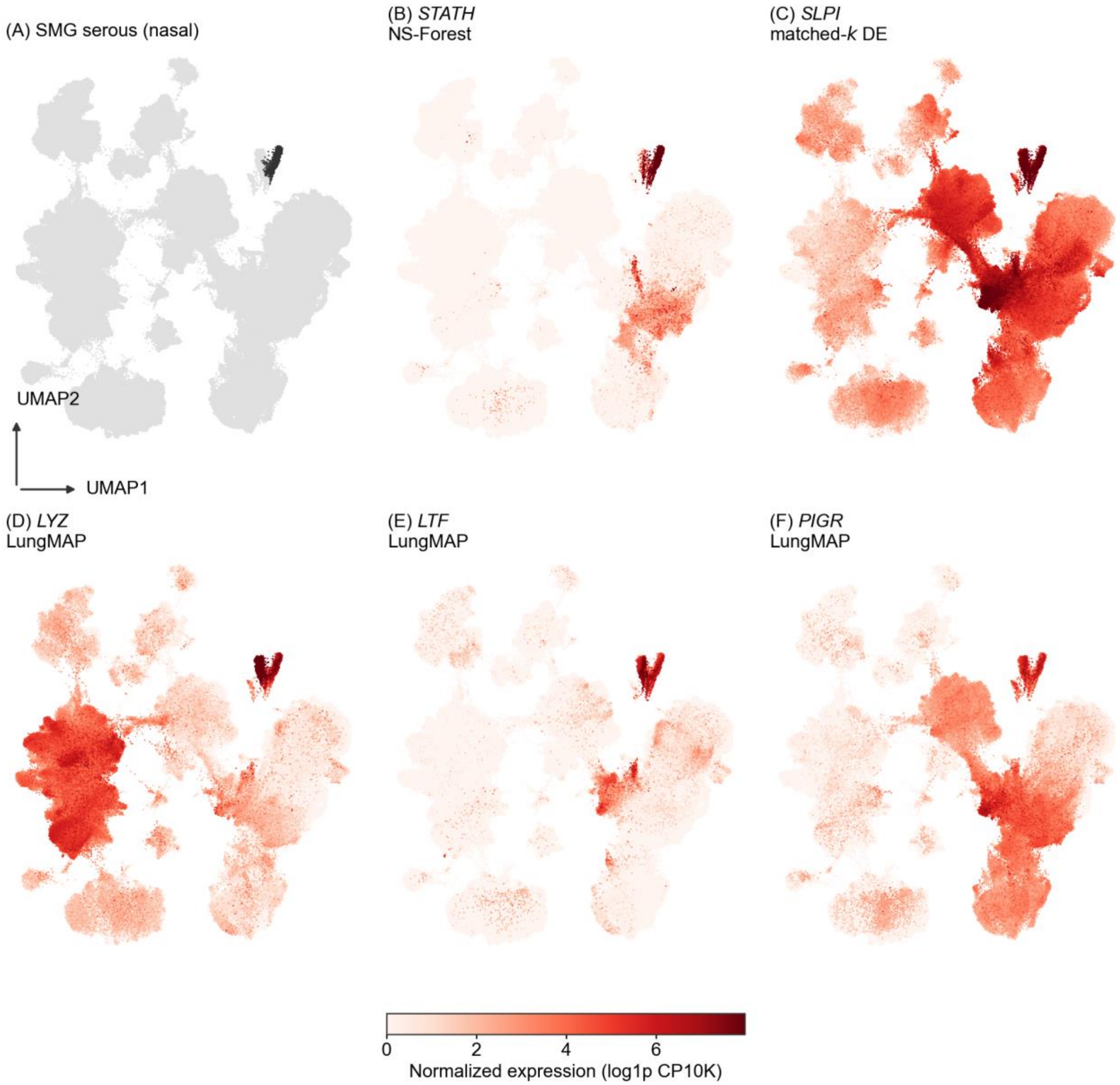


**Supplementary Figure S4. Location and marker gene expression of submucosal gland serous (nasal) cells on the HLCA Core UMAP. (A)** Location of target cluster (black) against all other cells (grey). **(B)** Expression profile of *STATH*, the NS-Forest marker. **(C)** Expression profile of *SLPI*, the top ranked DE marker. **(D-F)** Expression profile of *LYZ, LTF, PIGR*, the canonical markers listed by LungMAP cell cards (Sun et al., 2022). All expression panels share a color shade using normalized expression (log1p CP10K). *STATH* is largely confined to the target cluster while *SLPI* is detected broadly throughout the epithelial class.

**Supplementary Figure S5**

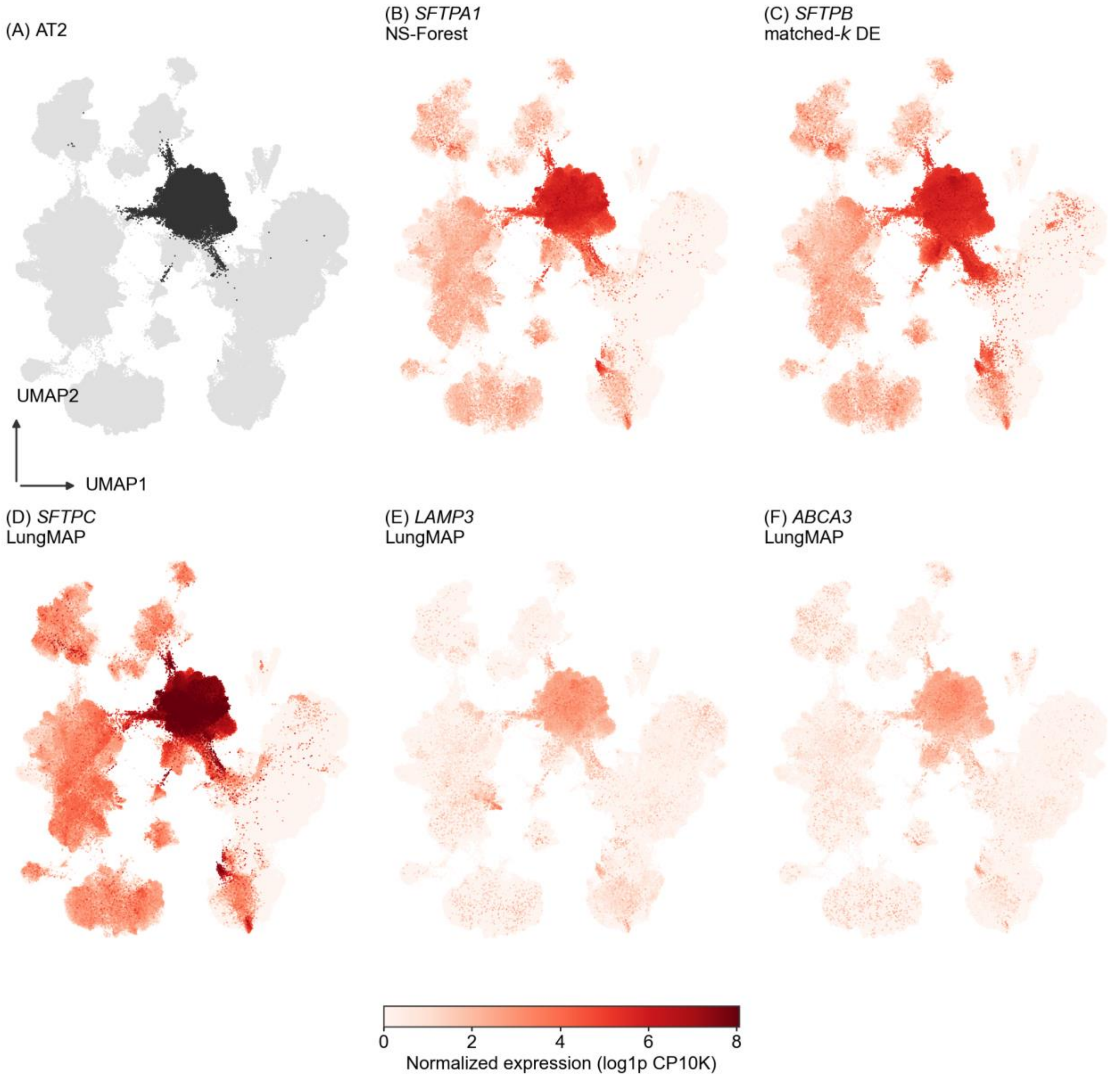


**Supplementary Figure S5. Location and marker gene expression of AT2 cells on the HLCA Core UMAP. (A)** Location of target cluster (black) against all other cells (grey). **(B)** Expression profile of *SFTPA1*, the NS-Forest marker. **(C)** Expression profile of *SFTPB*, the top ranked DE marker. **(D-F)** Expression profile of *SFTPC, LAMP3, ABCA3*, the canonical markers listed by LungMAP cell cards (Sun et al., 2022). All expression panels share a color shade using normalized expression (log1p CP10K).

**Supplementary Table**

**Supplementary Table S1**

| | HLCA ($n$=61) | | Kidney ($n$=76) | | Brain ($n$=75) | |
|---|---|---|---|---|---|---|
| Method | Pearson $r$ | Spearman $\rho$ | Pearson $r$ | Spearman $\rho$ | Pearson $r$ | Spearman $\rho$ |
| t-test | 0.68 | 0.927 | 0.77 | 0.906 | 0.78 | 0.796 |
| t-test-overest-var | 0.73 | 0.934 | 0.82 | 0.928 | 0.78 | 0.861 |
| logreg | 0.65 | 0.704 | 0.56 | 0.504 | 0.87 | 0.856 |
| **Wilcoxon** | **0.90** | **0.995** | **0.90** | **0.985** | **0.95** | **0.994** |
| **NS-Forest F-Score** | **0.10** | **0.127** | **0.28** | **0.264** | **0.21** | **0.209** |

**Supplementary Table S1. All methods tested against cluster size.** Correlations are computed against log10 cluster size on matched-k mean DE score for each cluster. One point was taken per cluster using the same procedure in Figure 1B with $n$ representing the number of clusters per dataset. Method names follow Scanpy's *rank_genes_groups* options with bolded rows marking the two methods used for analysis in the main text.
